# Rarγ -Foxa1 signaling promotes luminal identity in prostate progenitors and is disrupted in prostate cancer

**DOI:** 10.1101/2024.03.06.583256

**Authors:** Dario De Felice, Alessandro Alaimo, Davide Bressan, Sacha Genovesi, Elisa Marmocchi, Nicole Annesi, Giulia Beccaceci, Davide Dalfovo, Federico Cutrupi, Veronica Foletto, Marco Lorenzoni, Francesco Gandolfi, Srinivasaraghavan Kannan, Chandra S. Verma, Alessandro Vasciaveo, Michael M. Shen, Alessandro Romanel, Fulvio Chiacchiera, Francesco Cambuli, Andrea Lunardi

## Abstract

Retinoic acid (RA) signaling is a master regulator of vertebrate development with crucial roles in directing body axis orientation and tissue differentiation, including in the reproductive system. However, a mechanistic understanding of how RA signaling promotes cell lineage identity in different tissues is often missing.

Here, leveraging prostate organoid technology, we demonstrated that RA signaling orchestrates the commitment of adult mouse prostate progenitors to glandular identity, epithelial barrier integrity, and ultimately, proper specification of the prostatic lumen. Mechanistically, RA-dependent RARγ activation promotes the expression of the pioneer factor Foxa1, which synergizes with the androgen pathway for proper luminal expansion, cytoarchitecture and function. *FOXA1* nucleotide variants are common in human prostate and breast cancers and considered driver mutations, though their pathogenic mechanism is incompletely understood. Combining functional genetics experiments with structural modeling of FOXA1 folding and chromatin binding analyses, we discovered that FOXA1^F254E255^ is a loss-of-function mutation leading to compromised transcriptional function and lack of luminal fate commitment of prostate progenitors.

Overall, we define RA as a crucial instructive signal for glandular identity in adult prostate progenitors. We propose deregulation of vitamin A metabolism as a risk factor for benign and malignant prostate disease, and identified cancer associated FOXA1 indels affecting residue F254 as loss-of-function mutations promoting dedifferentiation of adult prostate progenitors.

Summary: Retinoic acid signaling orchestrates luminal differentiation of adult prostate progenitors

## Introduction

About forty years ago, androgen cycling experiments performed in rodents demonstrated the existence of stem/progenitor cell population(s) in the adult prostate epithelium capable of surviving castrate levels of testosterone (T) and retaining ample regenerative potential after androgens replenishment (*1*, *2*). However, the widespread cellular quiescence typical of the adult prostate epithelium has made extremely difficult the identification and characterization of such stem/progenitor cells and their niches. Early studies characterized cell populations expressing both basal and luminal markers, leading to the hypothesis of “*intermediate*” progenitor cells (*3*). Afterward, cell transplantation experiments and lineage tracing studies in animal models showed that both basal and luminal compartments contain multipotent progenitors characterized by extensive plasticity and capable of differentiating towards both luminal and basal cell lineages (*4–7*).

Recently, single-cell transcriptomics has greatly improved our knowledge of the different cell subtypes that populate the adult prostatic epithelium, formally defining the proximal anatomical district as the preferred niche for a variety of epithelial progenitor cells (basal, luminal proximal - LumP and periurethral - PrU), although they are also found in low abundance in the distal compartment (*8–13*). These progenitor cells, which are largely dormant in adult tissue (*14*, *15*), show great regenerative potential in *ex-vivo* assays (*8*, *14*). Lineage tracing experiments combined with genetic approaches in animal models have defined *NKX3.1*, *FOXA1*, *HOXB13*, *SOX9*, *AR* and *TRP63* genes as major players in prostate epithelium differentiation and morphogenesis (*16*). Among them, AR is undeniably the most studied and best characterized transcription factor acting in the prostate tissue.

The crucial role of endocrine and paracrine signals in prostate organogenesis is well-recognized. However, limited evidence connects specific signaling pathways with prostate epithelium differentiation and homeostasis (*17–19*). In 1993, Pierre Chambon described *RARγ*-null males as sterile due to squamous, instead of secretory, differentiation of the prostate epithelium and seminal vesicles (*20*), thus suggesting a critical function of retinoic signaling in the establishment of the luminal compartment of mouse prostate. Retinoids are vitamers of Vitamin A involved in many biochemical processes, including cell differentiation and embryonic development (*21*). The pleiotropic actions of retinoids are mediated by two families of nuclear receptors: retinoic acid receptors (RARs) α, β, and γ and retinoid X receptors (RXRs) (*22–24*). Upon RA binding, these receptors recognize specific DNA elements in the regulatory regions of selected genes and modulate their expression (*2*, *25*). Nevertheless, how RARγ transduces RA signaling into a transcriptional program that commits prostate progenitors to luminal fate, and then maintains the secretory epithelium, has remained unclear.

Here, leveraging organoid technology for the study of the prostate, we identify a new molecular link between retinoic acid (RA), its transcriptional mediator RARγ, and the pioneer transcription factor Foxa1, acting in prostate progenitors to enforce glandular identity in cooperation with androgen signaling and its nuclear receptor Ar. Considering that FOXA1 mutations are common in prostate cancer, we analyzed recurrent mutant isoforms and demonstrated that the most frequent coding alteration, FOXA1^F254E255^, is a hypomorphic variant with reduced chromatin binding, associated with progenitor dedifferentiation and loss of glandular identity. All-trans RA (ATRA) and RARγ agonists can boost FOXA1 expression and mitigate progenitor dedifferentiation. Our study paves the way for pharmacological strategies aimed to restore near-physiological FOXA1 activity in cancer cells.

## Results

### Retinoic acid promotes prostate-like cytoarchitecture and lumen formation in prostate organoids cooperating with androgen signaling

Organoid cultures were established from the prostate of inbred C57BL/6J and outbred CD1 mice based on the protocol originally described by Drost and colleagues (*26*). Briefly, small tissue fragments were enzymatically and mechanically digested, embedded into hydrogel droplets, and cultured in a defined medium including EGF, Noggin, R-spondin 1, the TGF-β inhibitor A83-01, and dihydrotestosterone (DHT) (hereafter ENRAD media, Figure 1A). Under such growth conditions, primary adult prostate cells proliferate rapidly forming 3D structures primarily made up of a compact and disorganized mass of cytokeratin 5 (Krt5)-positive basal cells intermixed with a small number of ectopically located cytokeratin 8 (Krt8)-positive cells (Figure 1B-C). Only a small fraction of the 3D structures showed a prostate-like cytoarchitecture displaying a double layer of basal and luminal cells correctly organized to form a lumen (hereafter mouse prostate organoids, mPrOs) (*18*, *19*) (Figure 1B-C).

**Figure 1.**
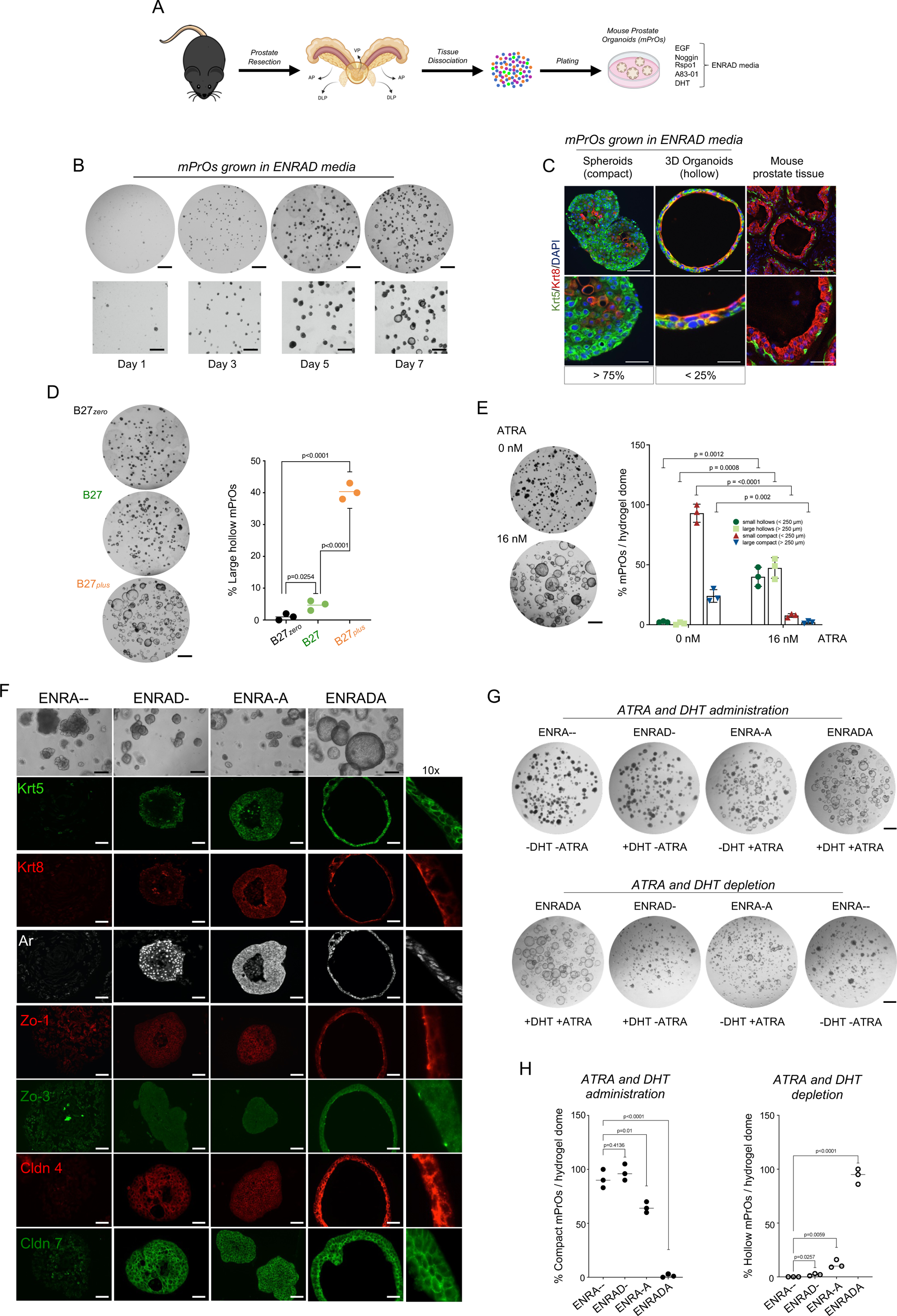
Retinoic acid promotes prostate-like cytoarchitecture and lumen formation in prostate organoids cooperating with androgen signaling. (A) Schematic overview of the procedure for establishing mPrOs (adapted from Karthaus et al. 2014). (B) Representative stereoscopic images of the mixed organoid population at different days of culture, scale bar: 1 mm. Magnifications (2x) are shown in the lower panels. Scale bars, 500 μm. N > 3 independent biological replicates. (C) Immunofluorescence staining of basal (Krt5) and luminal (Krt8) cytokeratins in mPrOs and mouse prostate tissue. Cell nuclei are stained with DAPI. Scale bars, mPrOs 50 μm; prostate 500 μm. Magnification of the selected area are shown. Scale bars, mPrOs 20 μm, prostate 200 μm. N > 3 independent biological replicates. (D) Representative stereoscopic images (left) and quantification (right) of mPrOs cultured in medium conditioned with different concentrations of ATRA (B27_zero_= 0 nM, B27 = 6 nM, B27_plus_ = 16 nM; a minimum of 100 organoids/dome x 3 domes were counted for each condition; large mPrOs, diameter >250 μm). Scale bar, 1 mm. N = 3 independent biological replicates. (E) Representative stereoscopic images (left) and quantitative phenotypic comparison (right) of mPrOs cultured with or without ATRA. Scale bar, 1 mm. N = 3 independent biological replicates. (F) Representative stereoscopic images and immunofluorescence analysis of Krt5, Krt8, Ar, Zo-1 (*Tjp1*), Zo-3 (*Tjp3*), Cldn4, and Cldn7 expression and localization in mPrOs cultured with or without DHT, ATRA or both. Scale bars, 100 μm; n = 2 independent biological replicates. Magnification (10x) of immunostaining of mPrOs cultured in presence of DHT and ATRA (ENRADA medium) are shown to pointing out protein localization. (G,H) Representative stereoscopic images (G) and quantitative analysis (H) of mPrOs morphology upon administration, or withdrawal, of DHT and ATRA. Scale bars, 1 mm. N = 3 independent biological replicates.

Considering the low efficiency in generating prostate-like organoids, we reviewed the literature and identified retinoic acid as a putative differentiation signal in the prostate, based on pioneering work by Pierre Chambon and colleagues on mouse genetic mutants (*20*). Nanomolar concentrations of retinol, a Vitamin A derivative that can be metabolized to retinoic acid (RA) (Supplementary Figure S1A), are present in the B-27 supplement (*27*) commonly employed for the growth of primary cells and organoids *in vitro* (*28*, *29*). The replacement of the standard B-27 supplement (hereafter B-27) with the vitamin A-free formulation (hereafter B-27 *zero*) resulted in the loss of the small percentage of large hollow mPrOs, leaving only compact spheroids in culture (Figure 1D). Conversely, administration of B-27 *zero* complemented with concentrations between 6 and 16 nM of all-trans retinoic acid (ATRA) (hereafter B-27 *plus* or ENRAD<u>A</u>) significantly boosted RA signaling (as demonstrated by activation of well-characterized target genes as *Aldh1a1* and *Rarb*) and enhanced the formation of mPrOs (Figure 1D-E, G-H and Supplementary Figure S1B-D) characterized by juxtaposed basal (Krt5) and luminal (Krt8) cells surrounding a well-defined expanded lumen (*18*, *19*) (Figure 1F). We observed proper polarization of luminal cells and epithelial barrier integrity, as visualized by immunostaining for apical (*e.g*., Zo1 and 3) and junctional markers (*e.g*., Cld4 and 7) (*30*) (Figure 1F). Notably, RA signaling was necessary but not sufficient to promote prostate-like cytoarchitecture and lumen formation in prostate organoids and acted in concert with the androgen pathway (Figure 1F-H and Supplementary Figure S1D). Analogous to androgen signaling (*18*), retinoic acid signaling displayed a fully reversible phenotypic switch in culture, as shown by cycling experiments (Figure 1G-H).

### A Rarγ-Foxa1 transcriptional cascade is essential for the retinoic acid control of luminal identity in adult prostate progenitors

To understand the molecular basis of the effect of retinoic acid signaling on prostate progenitors, we performed bulk RNA-seq on mPrOs grown in the absence or presence of DHT and ATRA ((ENRA-- (without ATRA and DHT) *vs*. ENRAD- (without ATRA with DHT) *vs.* ENRA-A (with ATRA without DHT) *vs.* ENRADA (with ATRA and DHT) conditions)). Comparing mPrOs growing with or without ATRA (e.g., ENRA-- and ENRAD- vs. ENRA-A and ENRADA, respectively), we identified *Foxa1,* the pioneer transcription factor of the luminal lineage, among the top differentially expressed genes activated by retinoic signaling (Figure 2A). In conjunction with *Foxa1* upregulation, we observed increased expression of the *Androgen Receptor* (*Ar*) and, conversely, downregulation of *Trp63*, which represent key transcription factors for the luminal and basal cell lineages, respectively (Figure 2D and Supplementary Figure S2A). Expanding our analysis to additional markers of basal and luminal cells, we found that ATRA prominently enhanced luminal markers, and especially luminal progenitor markers (*e.g*., *Krt4, Krt7, Clu, Wfdc2 and Ppp1r1b*) (*8*, *9*, *18*), with the highest levels observed upon supplementation of both DHT and ATRA (ENRADA) (Supplementary Figure S2A-B). Genes encoding for tight junctions’ proteins were also induced by retinoic acid (*e.g., Tjp1* and *Tjp3*, *Ocln*, *Cldn4* and *Cldn7*) (Supplementary Figure S2B). Conversely, the absence of ATRA (ENRA-- and ENRAD- in B27 *zero*) favored phenotypic and molecular features typical of stratified squamous epithelia, including the expression of late cornified envelope family genes (*e.g*., *Lce1e*, *Lc1f*, *Lc3d*, *Lcd3e* and *Lcd3f*) and the formation of spheroids almost entirely made up of basal cells enclosing anucleated cornified cells (Supplementary Figure S2C-D).

**Figure 2.**
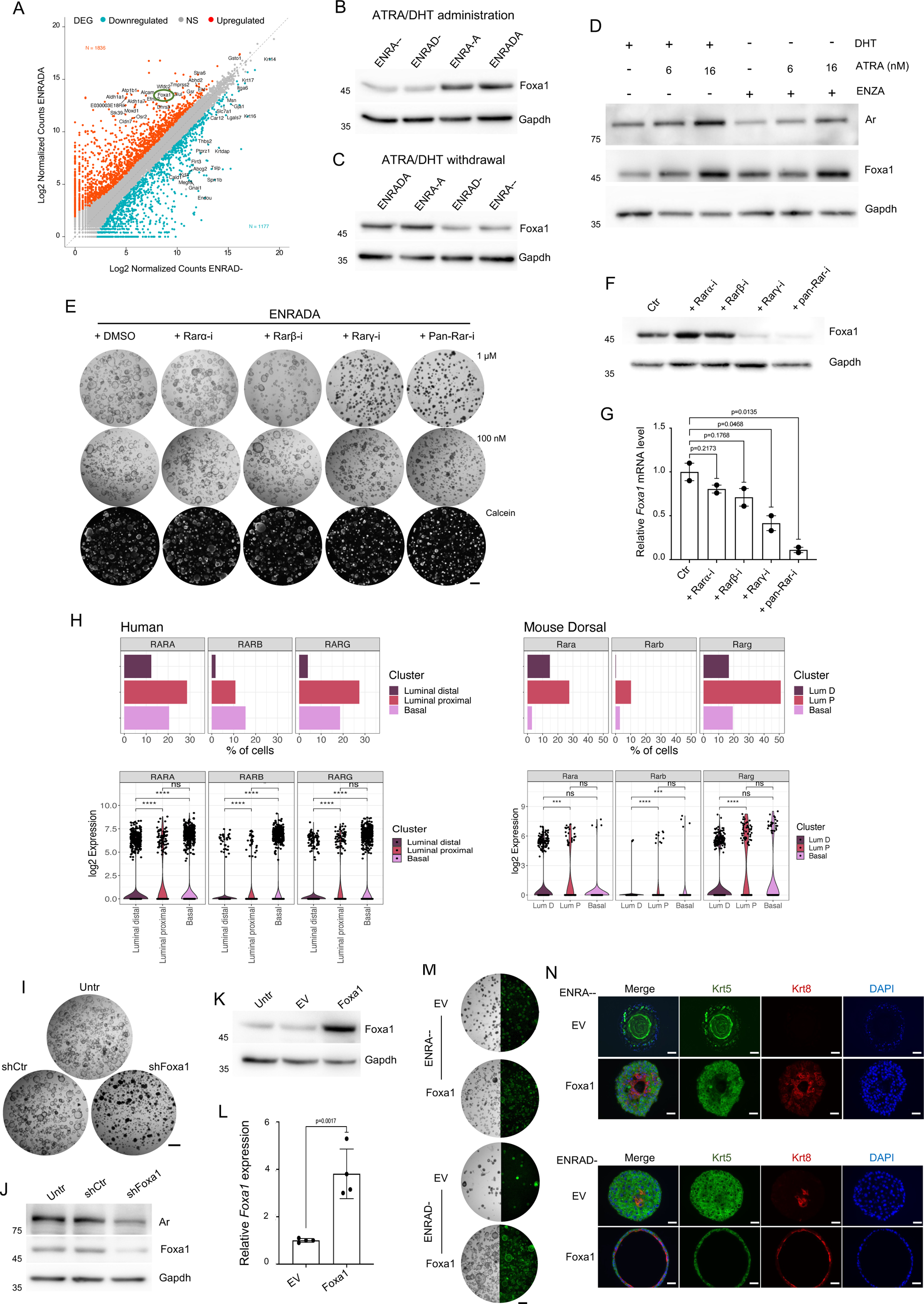
A Rarγ - Foxa1 transcriptional cascade is essential for the retinoic acid control of glandular identity in adult prostate progenitors. (A) Scatter plot representing the changes in gene expressions between enrad and enrada conditions. The number of significant up and down regulated genes is indicated as N in the figure. (B,C) Representative Western blot analysis of Foxa1 expression in mPrOs upon administration (B) and successive withdrawal (C) of ATRA and DHT individually or in combination. Gapdh is used as loading control. N = 3 independent biological replicates. (D) Representative Western blot analysis of Ar and Foxa1 expression in mPrOs upon administration or not of DHT (10 nM), ATRA (at different concentrations), and Enzalutamide (ENZA, 10 μM). Gapdh is used as loading control. N = 3 independent biological replicates. (E) mPrOs morphology upon administration of RARs inhibitors at different concentrations (upper and middle panels). Calcein staining determines mPrOs viability (lower panels). Scale bars, 1 mm. N = 3 independent biological replicates. (F) Representative Western blot analysis of Foxa1 expression in mPrOs treated with the different RAR inhibitors for six days. Gapdh is used as loading control. N = 3 independent biological replicates. (G) RT-qPCR analysis of Foxa1 expression in mPrOs treated with the RARs inhibitors. Data are presented as mean value ± s.d. of n = 2 independent biological replicates. (H) Percentage of cells (bar plots) and expression levels (violin plots) of, *RARα RARβ*, and *RARγ* genes in epithelial cell populations of human and mouse normal prostate (single cell data from Crowley et al., 2020). (I,J) Phenotypic response of mPrOs cultured in ENRADA to Foxa1 knock-down (I). Western blot showing reduction of Foxa1 and Ar level in mPrOs stably transduced with shRNAs against Foxa1 (J). Untransduced mPrOs and mPrOs expressing not targeting shRNAs (shCtr) are used as controls. Gapdh is used as loading control. N = 3 independent biological replicates. (K,L) Western blot analysis of Foxa1 expression in mPrOs grown without DHT and ATRA (ENRA--) untransduced (Untr), stably transduced with an empty vector (EV) or with a vector expressing mouse Foxa1 (Foxa1) (K). Gapdh is used as loading control. RT-qPCR analysis of Foxa1 RNA expression in EV and Foxa1 mPrOs cultured in ENRA-- medium (L). N > 3 independent biological replicates. (M,N) Morphological comparison of wild-type and transduced (EV and Foxa1) mPrOs cultured without ATRA and with or without DHT (ENRAD-, ENRA--) (M). Scale bar, 1 mm. N > 3 independent biological replicates. Immunofluorescence analysis of Krt5 (basal) and Krt8 (luminal) markers in the different conditions. Nuclei are stained with DAPI (N). Scale bar 50 μm.

Retinoic acid supplementation markedly increased Foxa1 protein expression, which was independent from androgen signaling and Ar transcriptional activity (as demonstrated by enzalutamide treatment) (Figure 2B-D and Supplementary Figure S3A-B). To gain mechanistic insights into the control of prostate organoid cytoarchitecture and Foxa1 expression by retinoic acid signaling, we targeted retinoic acid receptors (Rar*s*) with isoform-specific or a pan-Rar inhibitors. We found that RARγ inhibition (RARγ-i) significantly reduced lumen formation as well as Foxa1 mRNA and protein levels in prostate organoids (Figure 2E-G). Leveraging publicly available datasets (*8*), we mapped *RARγ* expression in the normal mouse and human prostate epithelium at the single-cell level. We found that *RAR*γ is predominantly expressed by progenitor cells *in vivo*, being highly transcribed by nearly 20% of basal and 30% of luminal proximal progenitors in human, and by 20% of basal, 50% of LumP and 40% of PrU in the mouse (Figure 2H and Supplementary Figure S3C). Crucially, Foxa1 is essential for mediating the control of retinoic acid on glandular identity in prostate progenitors. Foxa1 knock-down abolished the ability of RA to generate prostate organoids that have a luminal cavity (Figure 2I-J). Conversely, constitutive expression of a transgene encoding for Foxa1 leads to luminal priming even in the absence of ATRA and DHT (ENRA-- conditions) and to the formation of large hollow mPrOs with a well-structured prostate-like luminal compartment if androgens are added (ENRAD-) to retinoic-depleted media (Figure 2K-N and Supplementary Figure S3C-D).

### Foxa1 occupies enhancers and promoters of key luminal progenitor genes and reshapes genome-wide androgen receptor binding

To shed light on how Foxa1 and Ar transcription factors coordinately promote a luminal progenitor gene expression program and a glandular phenotype in prostate progenitors, we combined our RNA-seq analysis of mouse prostate organoids (mPrOs) grown with or without ATRA and DHT with publicly available Foxa1 and Ar ChIP-seq datasets from mPrOs constitutively expressing a Foxa1 transgene or a control empty vector (EV) (*31*) (Figure 3A).

**Figure 3.**
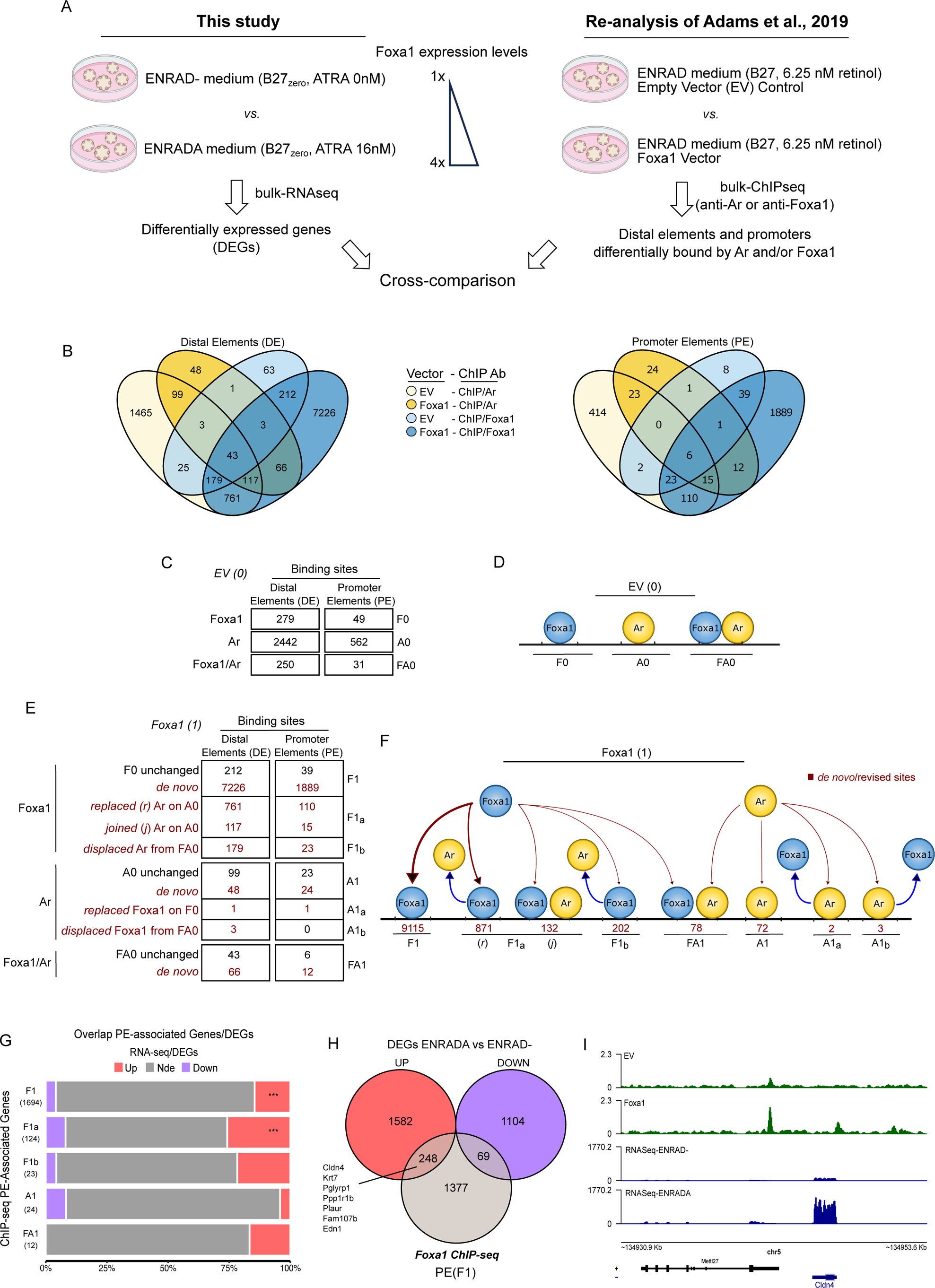
Foxa1 occupies distal and promoter elements of key luminal progenitor genes and reshape androgen receptor binding genome-wide. (A) Schematic representation of the cross-comparison study of RNA-seq analysis performed in this study and published ChIP-seq datasets (Adams et al., 2019). (B) Venn diagram showing the binding sites of Ar and Foxa1 in the genome of mPrOs expressing endogenous (empty vector, EV) or exogenous (Foxa1) Foxa1. Distal elements (± 2.5 kb away from an annotated gene promoter) are displayed on the left, whereas promoter sites are shown on the right. ChIP-seq data are from Adams et al. 2019 (n = 2 replicates per condition). (C-F) Numerical (C and E) and graphical (D and F) representation of Foxa1 and Ar cistromes at both distal (DE) and promoter (PE) elements with endogenous levels of Foxa1 (EV (0)) and upon its over-expression (Foxa1 (1)). The number of *de novo* and pre-existing but rearranged PE/DE sites are indicated in red. (G) Percentage overlap of ChIP-seq promoter elements (PE)-associated Genes (Adams et al., 2019) with Upregulated and Downregulated DEGs from RNA-seq (this work). The significance of the overlap has been determined by a hypergeometric test and denoted by asterisks. Each bar represents a set of genes (*e.g.,* n = 1694) associated to a specific class of PE (*e.g.* de novo Foxa1-bound PE, F1). (H) Venn diagrams showing the overlap of differentially expressed genes in mPrOs cultured in ENRADA versus ENRAD, and exogenous Foxa1-bound promoter elements (PE (F1)) in the genome. Relevant up-regulated genes in the intersection are highlighted. (I) Genomic snapshot of ChIP-seq (n = 2 pooled replicates) and RNA-seq (n = 3 pooled replicates) signals over the selected gene *Cldn4*.

In standard organoid culture conditions, RA signaling is limited (retinol < 10 nM) and Foxa1 levels are low. Transgenic expression of a lentiviral vector encoding for *Foxa1* mimics ATRA treatment leading to roughly a four-fold increase in Foxa1 expression in comparison to the empty vector control (*31*) (Figure 2K-L). Reanalysis of Foxa1 ChIP-seq in mPrOs^Foxa1^ *vs*. mPrOs^EV^ led to the detection of more than 7,000 Foxa1-bound (F1) distal regulatory elements (F1-DE) and almost 2,000 promoters (F1-PE), consistent with the known pioneer ability of this transcription factor (*32*) (Figure 3B-F and Supplementary Table S1). The intersection of the ChIP-seq datasets for Foxa1 with the list of genes that increase in expression in organoids treated with ATRA (ENRADA) compared to those kept in regular medium (ENRAD-) (Supplementary Table S2) highlighted key luminal and intermediate prostate progenitor markers (*e.g*., *Krt7*, *Ppp1r1b, Plaur*) (*8*, *9*, *18*), luminal lineage transcription factors (e.g. *Foxa1, Nkx3.1*) (*16*), genes involved in epithelial barrier establishment (*e.g*., *Cldn4, Tjp3, Tjp1*) (*30*) and luminal cells function (*e.g*., *Trpm8, Krt7, Krt8*) (*8*, *9*, *18*, *33–35*) among the principal targets of Foxa1 transcriptional activity (Figure 3B-I, Supplementary Figure S4A-B and Supplementary Table S3).

In addition to the primary activity of Foxa1 on crucial epithelial genes, our analysis revealed widespread Foxa1-mediated reprogramming of Ar. In mPrOs^EV^, Ar occupied over 3,000 genomic sites, including ∼ 2,500 putative enhancers and > 500 gene promoters (Figure 3B-F, Supplementary Figure S4C-H, and Supplementary Table S1). Foxa1 expression in mPrOs^Foxa1^ was associated with the extensive reprogramming of the Ar cistrome. In mPrOs^Foxa1^, Foxa1 replaced Ar at ∼ 35% of the putative enhancers (distal elements, DE; Ar-bound (A) DE in mPrOs^EV^ (0), A0 n=2442; A0 where Ar is replaced *(r)* by Foxa1 (F) in mPrOs^Foxa1^ (1), F1_a_ *(r)* n=761) and at ∼ 20% of the promoters (Promoter elements, PE; A0 n=562, F1_a_ (*r)* n=110) bound exclusively by Ar in mPrOs^EV^. Finally, ∼ 70% of the genomic loci occupied by both transcription factors in mPrOs^EV^ (DE+PE; FA0 n=250+31) were exclusively occupied by Foxa1 in mPrOs^Foxa1^ (DE+PE; F1b n=179+23) (Figure 3C-F). In the group of loci occupied by Ar in mPrOs^EV^ and in which Foxa1 replaced Ar in mPrOs^Foxa1^ (F1_a_ (*r*)), we found the distal elements (DE) of the progenitor markers *Wfdc2*, *Krt19*, *Sox5*, *Fgf1* and *Runx2* and the luminal-associated genes *Krt8*, *Cldn4* and *Steap4* (*8*, *9*, *11*, *13*, *18*, *36*), which result upregulated upon ATRA treatment (Figure 3C-F, Supplementary Figure S4C and Supplementary Table S3). Additional critical DEGs, such as the key progenitor marker *Clu* (*8*, *9*) and *Il33*, a cytokine involved in epigenetic reprogramming in epithelia (*37*), displayed Foxa1/Ar replacement (F1_a_ (*r*)) at genes’ promoter elements (PE) (Figure 3C-F, Supplementary Figure S4D and Supplementary Table S3). Unexpectedly, the intersection of ChIP-seq (A1+A1_a_+A1_b_ and F1_a_ (*j*) + FA1, PE and DE) and RNA-seq analyses did not provide clear insights into the role of Ar in epithelial differentiation and lumenogenesis (Figure 3C-F, Supplementary Figure S4E-H and Supplementary Table S3). However, functional enrichment analysis of DEGs revealed the GO terms ‘Epithelial Cell Differentiation’ and ‘Tissue Remodeling’ as the two most highly enriched terms upon ATRA treatment (Supplementary Table S4).

### The hotspot Foxa1^F254E255^ prostate cancer mutant is impaired in promoting luminal identity in prostate progenitors

*FOXA1* is altered in ∼12% of prostate cancer patients, predominantly through single-residue variants and short indels (∼8.5% of cases). Among these mutations, about half (∼ 4.25%) occur within the Wing2 region (between H247 and E269) of the Forkhead DNA-binding domain (FKHD), which can be considered a mutational hotspot (*31*). A previous systematic phenotypic and molecular analysis of FOXA1 mutants in prostate organoids concluded that Wing2 alterations are gain-of-function variants conferring an enhanced pro-luminal differentiation program (*31*). Under standard organoid culture conditions, limited RA signaling results in low levels of Foxa1 and only a small fraction of prostate progenitors acquires luminal identity. We thus set out to investigate the role of the most common FOXA1 mutations in our optimized prostate organoid model, which is characterized by efficient luminal differentiation. In-frame indels are a common type of cancer mutations affecting *FOXA1* (*31*, *38*, *39*), and F254E255 is one of the most frequent variants affecting prostate cancer patients (Figure 4A).

**Figure 4.**
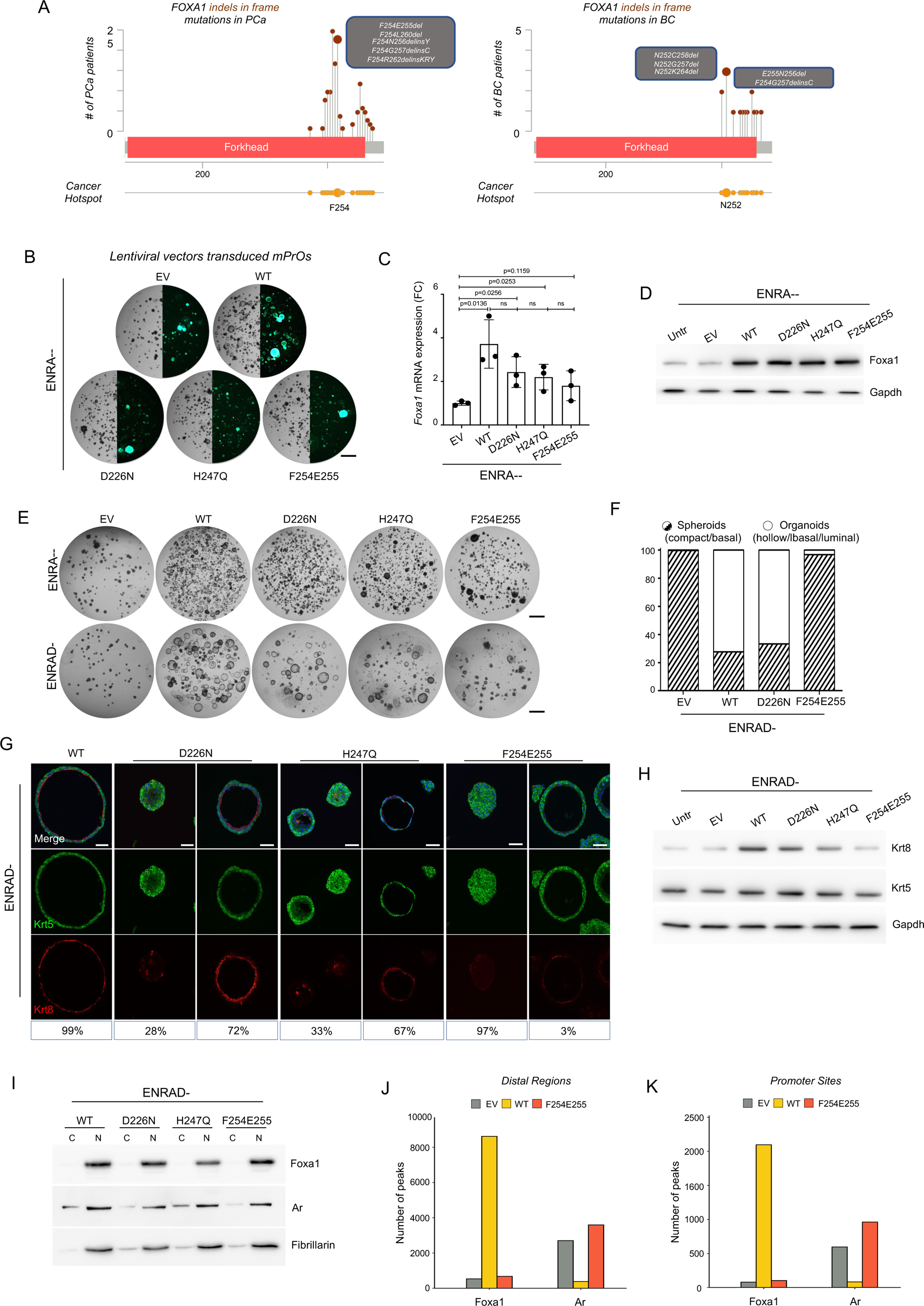
The hotspot Foxa1^F254E255^ prostate cancer mutation is unable to promote luminal identity in prostate progenitors. (A) Indels mutations of the alpha-helix region at the C-terminal part of the Forkhead domain of FOXA1 identified in in prostate (left) and breast (right) cancers (cBioportal/Cosmic databases). (B-D) Brightfield and fluorescence images of mPrOs grown without ATRA and DHT (ENRA--) expressing wild-type Foxa1 or its mutant forms D226N, H247Q, or F254E255. mPrOs transduced with the empty vector (EV) were used as controls (B). Scale bar, 1 mm. RT-qPCR analysis of Foxa1 RNA expression in transduced mPrOs cultured in ENRA-- medium (C). Western blot analysis of Foxa1 protein levels in transduced mPrOs (D). Gapdh is used as loading control. N = 3 independent biological replicates. (E,F) Morphological analysis of mPrOs transduced with Foxa1 variants upon re-administration of DHT (E) and quantification of compact versus hollow organoids (F). Scale bars: 1 mm. N = 2 independent biological replicates. (G,H) Immunofluorescence (G) and Western blot (H) analyses of Krt5 (basal) and Krt8 (luminal) markers in mPrOs cultured in presence of DHT but not ATRA (ENRAD-) and expressing exogenous wild-type Foxa1 or its mutant forms. Gapdh is used as loading control. Scale bar, 100 μm. N = 3 independent biological replicates. (I) Biochemical fractionation of nuclear (N) and cytosolic (C) compartments showing nuclear localization of wild type and mutant form of Foxa1. Ar and Fibrillarin are used as nuclear markers and loading controls. (J,K) Number of peaks identified by ChIP-seq in distal regions (J) and gene promoters (K) for Foxa1 and Ar in mPrOs stably transduced with wild-type Foxa1 (WT), Foxa1^F254E255^ (F254E255) or the empty vector (EV) and cultured with DHT but not ATRA, as reported in Adams et al., 2019.

We generated mouse prostate progenitor lines stably expressing F254E255 or two FOXA1 missense mutations occurring in the FKHD domain either before (D226N) or within the Wing2 region (H247Q). Organoids transduced with the empty vector (mPrOs^EV^) served as control (Figure 4B and Supplementary Figure S5A). All three Foxa1 mutants were expressed at similar levels than exogenous Foxa1 wild type (Foxa1^wt^), displaying a three- to four-fold increase in comparison to untransduced and mPrOs^EV^ control organoids (Figure 4C-D). In the absence of DHT and RA signaling (ENRA-- conditions with B27 *zero*), control organoids (mPrOs^EV^) rarely showed well-shaped hollow organoids (Figure 4B,E). As expected, the frequency of well-shaped hollow organoids increased in those expressing the wild-type form of Foxa1 (mPrOs^WT^), while the three *Foxa1* mutant organoid lines mainly generated compact spheroids, similar to controls (mPrOs^EV^) (Figure 4B,E). DHT administration (ENRAD-conditions) rescued the ability of mPrOS^WT^, mPrOS^D226N^ and mPrOs^H247Q^ to form hollow organoids, whereas the frequency remained <5% in mPrOs^F254E255^ (Figure 4E-G). Immunofluorescence and Western blotting experiments for prostate basal (Krt5) and luminal (Krt8) cell markers suggested a slight decrease in the ability of Foxa1 mutants D226N and H247Q to promote differentiation and expansion of luminal progenitors, a phenotype that was more severe for the F254E255 mutant (Figure 4G-H). Despite unperturbed nuclear localization of wild-type and mutants Foxa1 (Figure 4I and Supplementary S5B), reanalysis of the publicly available Foxa1 and Ar ChIP datasets in prostate organoids (*31*) revealed low occupancy of Foxa1^F254E255^ at distal and proximal DNA elements (Figure 4J-K and Supplementary Figure S5C). Peak numbers of Foxa1^F254E255^ were markedly distinct from wild-type Foxa1 and comparable to the EV control, as was its ability to displace Ar from DE and PE genome-wide (Figure 4J-K and Supplementary Figure S5C).

### Molecular modeling of the FOXA1^F254E255^ FKHD domain is consistent with impaired DNA binding ability

To gain a structural understanding of the impact of FOXA1 FKHD domain variants on DNA binding we computationally modeled such interactions (Figure 5). Molecular dynamics simulations (aMD) of wild-type FOXA1 bound to DNA yielded a stable complex, with most sampled conformations remaining around ∼3 to 4 Å from the original crystallographic structure for both interacting macromolecules (*e.g*., FOXA1 and the double-stranded DNA molecule) (Figure 5A, left panel-black line). The double-stranded DNA within the FOXA1^F254E255^-DNA complex was also stable with sampled conformations within ∼4 Å of the reference structure (Figure 5A, left panel-red line). However, compared to wild-type FOXA1 (Figure 5A, right panel-black line), mutant FOXA1^F254E255^ deviated significantly from its initial conformation, averaging ∼7 Å (Figure 5A, right panel-red line). The F254 and E255 residues deleted in mutant FOXA1^F254E255^ are part of an α-helix at the C-terminus of the Forkhead domain (FKHD). The α-helix remained very stable during the simulation of wild-type FOXA1 binding to DNA (Figure 5B, D (black line)), whereas it completely unfolded in mutant FOXA1^F254E255^ (Figure 5C-D (red line)). Residue F254 is buried in a cavity formed by residues F72, I20, L90, C71, T24 and Y103, while the side chain of E255 is involved in hydrogen bond interactions with K189 and G184 (Figure 5B). These interactions were well-maintained during the MD simulation with wild-type FOXA1, whereas they were completely lost in FOXA1^F254E255^, leading to the unfolding of the α-helix at the C-terminus of the FKHD domain resulting in high flexibility. These structural changes in mutant FOXA1^F254E255^ are consistent with a reduction in the number of FOXA1-DNA contacts, causing loss of DNA affinity (Figure 5E).

**Figure 5.**
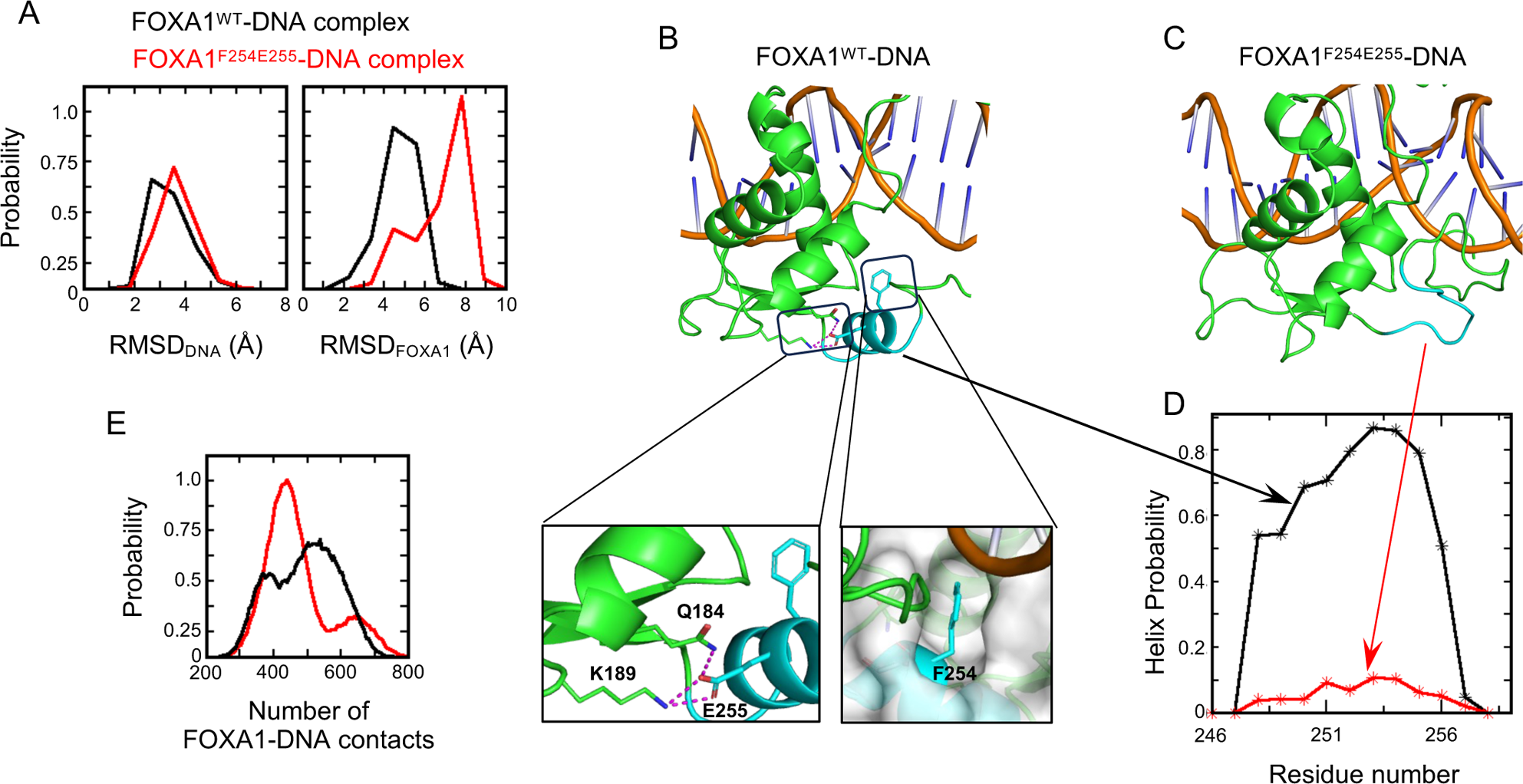
Molecular modeling of the FOXA1^F254E255^ FKHD domain is consistent with impaired DNA binding ability. (A) Distribution of Root Mean Square Deviation of conformations of FOXA1^WT^–DNA (black) and FOXA1^F254E255^–DNA (red) sampled during aMD simulations. (B,C) MD snapshot (from the MD simulations) highlighting the C-terminal α-helix (cyan) of FKHD of the FOXA1^WT^–DNA complex (B), which becomes disordered (cyan) in the FOXA1^F254E255^–DNA complex (C). In the case of the FOXA1^WT^–DNA complex (B), residues Q184, K189 and E255 are shown as sticks and interactions between them are highlighted in dashed lines (magenta). The residue F254 is shown as sticks and the region around it is shown as surface (grey) highlighting the burial of the sidechain of F254 in the cavity. (D) Distribution of the helix probability (from the MD simulations) of the conformation of the C-terminal α-helix of FKHD of FOXA1 sampled during FOXA1^WT^–DNA (black) and FOXA1^F254E255^–DNA (red) complexes. (E) Probability of the number of FOXA1–DNA contacts of FOXA1^WT^ (black) and FOXA1^F254E255^ (red) (from the MD simulations).

## Discussion

Vitamin A (retinol) is a nutrient with well-recognized critical roles in embryonic development, organogenesis, tissue differentiation, and homeostasis. Since its discovery a century ago, developmental defects associated with either an excess or a deficit of vitamin A (VAD syndrome) have led to a profound interest in defining the mechanisms of action that might explain its pleiotropy.

Identification of the active derivative of Vitamin A, the retinoic acid (RA), and its receptors (RARα (*40*), β (*41*), and γ (*42*, *43*)) paved the way for the molecular and functional characterization of RA signaling. RARα is widely expressed in almost all organs and tissues of the body while RARβ and RARγ show more restricted patterns (*21*). Genetic knockout of the *Rarγ* gene in mice unveiled the primarily role of this receptor in spermatogenesis (*44*) and development of the secretory epithelium of seminal vesicles and prostate (*20*). However, the molecular circuits underlying these processes have been understudied due to the lack of accessible models for precise cellular and molecular investigations.

Our work on mouse prostate organoids has shed light on *Foxa1* as a crucial target of RA-RARγ signaling in the prostate. In rodent development, *Foxa1* is expressed in the entire urogenital epithelium (UGE) before prostate induction, while it is restricted to the luminal compartment thereafter. *Foxa1* genetic ablation causes loss of luminal secretory cells, prostatic hyperplasia and expression of seminal vesicles markers, all phenotypes consistent with a crucial role of Foxa1 in luminal lineage differentiation (*45*, *46*). Consistent with these findings, we have shown that nanomolar amounts of all-trans retinoic acid (ATRA) induce the expression of mouse prostate luminal progenitor genes (*Krt4, Krt7, Wfdc2, Clu, Ppp1r1b*) as well as genes *(Tjp1, Tjp3, Ocln, Cldn4, Cldn7)* encoding tight- and gap-junction proteins necessary to establish a precise, solid, and functional cytoarchitecture of the luminal progenitors compartment. Mechanistically, the pioneering activity of Foxa1 in the prostate epithelium has been attributed to its ability to shape androgen signaling by favoring Ar activity on promoter and enhancer regions of specific subsets of genes (*31*, *47–49*). Yet, we found that Foxa1 binds the distal and/or promoter regions of many genes independently of Ar, suggesting direct and central transcriptional control of the luminal progenitor fate. Still, the androgen pathway is required for proper lumenogenesis, and thus for the formation of well-shaped hollow organoids.

While our work greatly advances the ability to model the luminal compartment *in vitro*, organoid systems currently lack robust and reproducible secretion of kallikrein-rich fluids typical of terminal differentiated luminal cells. We expect that progress in recapitulating physiological prostatic functions *in vitro* will further extend our comprehension of the underlying molecular mechanisms, including the interplay between FOXA1 and AR transcriptional regulation. Notably, while complete withdrawal of RA signaling from the growth medium generates compact spheres of proliferating cells that are almost invariably positive for the basal marker Krt5, we did rarely observe a few ectopic Krt8 positive cells in the center of the Krt5+ mass of cells, pointing to the possible role of still unknown signaling mediators in the specification of luminal progenitors.

The identification of Foxa1 as a major target of RA signaling led us to explore its role in prostate cancer. Recently, recurrent missense mutations have been identified in human prostate and breast cancers as drivers of epithelial transformation and tumorigenesis (*31*, *38*). FOXA1 mutations have been generally hypothesized as enhancers of transcriptional activity on canonical and *de novo* target genes, and as causal agents of aberrant androgen and estrogen receptor functions (*31*, *38*, *39*). Notably, the F254 residue, which plays a crucial role in stabilizing the α-helix at the C-terminus of the FKHD domain of FOXA1, is deleted in a large fraction of indel mutations in prostate and breast cancer (Figure 4A). We have shown that Foxa1^F254E255^ mutant fails in promoting the luminal lineage in mouse prostate organoids due to reduced DNA binding stability and impaired transcriptional activity.

FOXA1 loss-of-function mutations could represent a relevant genetic condition in oncology. FOXA1 binds DNA as a monomer on A(A/T)TRTT(G/T)R(C/T)T(C/T) consensus elements, or as a homodimer on compact palindromic DNA elements (diverging half-sites-DIV) (*50*, *51*). In cancer cells with loss-of-function mutations of FOXA1 that preserve the ability to form homodimers, concomitant induction of both wild-type and mutant alleles will presumably result in a dominant negative effect of the mutant protein on the regulation of DIV elements. In contrast, a benefit of FOXA1 overexpression should be expected on DIV controlled genes in the presence of loss-of-function mutant alleles unable to homodimerize, and on targeted genes where FOXA1 works as a monomer.

Whether and how this specific impairment of FOXA1 transcriptional function may impact tumor prognosis and treatment is still unknown but it deserves close attention for its potential clinical relevance. ATRA and synthetic retinoids such as fenretinide (4-HPR) or etretinate (Tegison) have been clinically tested in several solid tumors characterized by dysfunctional RA signaling (*52*, *53*). To date, no clinical trial has demonstrated efficacy, and activation of the retinoic pathway remains a clinical option only for the treatment of PML-RARα Acute Promyelocytic Leukemia (APL). Noteworthy, subgroups of patients with superficial papillary or resected high-risk non-muscle invasive bladder cancer showed reduced recurrence rate and cancer progression (*54–56*), while few patients with advanced breast cancer achieved partial response or had stable disease (*57*).

Overall, our study adds new important insights to the network of signaling pathways and molecular circuits regulating prostate progenitor commitment and adult tissue homeostasis and paves the way for more accurate assessments of retinoid derivatives for the treatment of specific forms of solid tumors that can be stratified by FOXA1 mutations.

## Materials and Methods

### Mouse Housing and Husbandry

Housing systems followed FELASA guidelines and recommendations concerning animal welfare, health monitoring and veterinary care, in compliance with the Directive 2010/63/UE and its Italian transposition D. L.vo 26/2014. Mice were monitored daily for general health and well-being and sentinel mice are used for quarterly monitoring for specific pathogens. Wild-type C57BL/6J (JAX # 000664) mice were purchased from the Jackson Laboratory. Mice were housed in room with 21 °C temperature with 12 hours light/dark cycle with light gradually rising at 7:00 a.m. and gradually decreasing at 7.00 p.m. A maximum of 5 animals were accommodated in IVC cages with food and water ad libitum and nesting materials and cardboard tunnels were provided as enrichment. Animal were sacrificed according to the European Communities Council Directive (2010/63/EU) and following the protocol approved by the Italian Ministry of Health and the University of Trento Animal Welfare Committee (642/2017-PR).

### Mouse prostate organoid cultures

Mouse prostate organoids were generated from prostate glands collected from adult (6 month-old) inbred C57BL/6J or outbred CD1 wild type males as described in (*18*). The following media were used (see Appendix Table 1 for small molecules used for organoid culture media):

- ENRADA: AdDMEM 4+, 50 ng/ml Egf, 100 ng/ml Noggin, 10% R-Spondin1 conditioned medium (Cell Tech Facility at CIBIO, using Cultrex® Rspo1 cells following the guidelines from Trevigen), 0.2 μM A83-01, 10 nM DHT, 16 nM ATRA. Stored in the dark at 4°C for up to one week.
- ENRAD-: AdDMEM 4+, 50 ng/ml Egf, 100 ng/ml Noggin, 10% R-Spondin1 conditioned medium (Cell Tech Facility at CIBIO, using Cultrex® Rspo1 cells following the guidelines from Trevigen), 0.2 μM A83-01, 10 nM DHT. Stored in the dark at 4°C for up to one week.
- ENRA--: AdDMEM 4+, 50 ng/ml Egf, 100 ng/ml Noggin, 10% R-Spondin1 conditioned medium (Cell Tech Facility at CIBIO, using Cultrex® Rspo1 cells following the guidelines from Trevigen), 0.2 μM A83-01. Stored at 4°C for up to one week.

### Viral transduction

Organoids were dissociated to single cells, and approximately 50,000 cells used for viral transduction. Spinoculation was performed in a low-adhesion 96 well-plate using 0.6-1.0 RTU/ml of lentiviral solution, supplemented with polybrene (4 μg/mL; Sigma Aldrich, H9268) and ENRADA complete medium (Egf (50 ng/mL; PeproTech, 315-09), Noggin (100 ng/mL; PeproTech, 120-10C), R-Spondin1 (10% conditioned medium), A83-01 (200 nM; Tocris, 2393), dihydrotestosterone (10 nM; Merck, 10300), and ATRA (16 nM; Merck, R2625)), or ENRA-- medium (without dihydrotestosterone and ATRA) supplemented with Y-27632 (10 μM; Calbiochem, 146986-50-7)) to a final volume of 300 μL. The plate was centrifuged for 1 hour at 600 g, cells resuspended in 200 μL of ENRADA or ENRA-- medium supplemented with Y-27632 (10 μM) and incubated in suspension at 37 °C for 4-6 hour. Then cells were mildly centrifuged (300 g, 5 min), cell pellet resuspended in 80% growth factor-reduced basement matrix (either Matrigel®, Corning, 356231; or BME-2®, AMSBIO, 3533) and seeded at the concentration of approximately 50,000 cells/mL by depositing at least six 40 μL drops at the bottom of a non-tissue culture treated plate. Domes were left to solidify for 15 minutes and covered with ENRADA or ENRA-- medium. Antibiotic selection started two days post-transduction. After approximately two weeks of antibiotic selection, transduced organoids expressed constitutively the green fluorescent protein (GFP). The following plasmids were used: pMSCV-Neo-GFP/FOXA1 (Addgene #105506) plasmid and the negative control pMSCV-Neo-GFP/Empty (Addgene #105505) were purchased on Addgene. FOXA1 mutated cDNAs (Twist Biosciences) were subcloned into pMSCV-Neo-GFP/FOXA1 after the enzymatic removal of the wild type FOXA1 cDNA to generate pMSCV-Neo-GFP/FOXA1^D226N^, pMSCV-Neo-GFP/FOXA1^H247Q^, and pMSCV-Neo-GFP/FOXA1^F254E255^. LEPG-shFoxa1 2959 and LEPG-shRLuc were kindly provided by Cristopher Vakoc’s Lab (*58*).

### Immunofluorescence studies

Organoids were cultured for 5-7 days, released from the basement membrane using a recovery solution – including Dispase II (1 mg/mL) – seeded in a neutralized collagen type-I solution (Corning, 354249) and cultured for additional 24 hours before fixing them with 4% paraformaldehyde (Sigma Aldrich, P6148) for 5 hours, at room temperature. Prostate tissue was harvested and immediately fixed using the same conditions. Paraffin embedding and 5 µm sectioning were carried out according to standard procedures. For immunolocalization studies, antigen retrieval was performed with citrate-based buffer (pH 6.0) (Vector Lab, H3300) in a microwave (90 - 100°C) for 20 minutes. Slides were incubated in blocking solution (5% FBS + 0.1% Triton-X in PBS) for 1 hour at room temperature, and with primary antibodies at 4 °C overnight. Spectrally distinct fluorochrome-conjugated antibodies were incubated for 2 hours at room temperature. Slides were counterstained with Hoechst 33342 (Abcam, ab145597), and FluorSave mounting medium (Merck, 345789) applied before the coverslip. Mouse prostate was isolated, fixed in 4% paraformaldehyde for 20 minutes at room temperature and processed for immunolocalization studies as described for organoids. Primary and secondary antibodies used in this study are listed in the Appendix Table 2.

### RNA extraction

Total RNA was extracted using the RNeasy Plus Micro kit (Qiagen, 74034) according to the manufacturer instructions, and analyzed with an Agilent BioAnalyzer 2100 to confirm integrity (RIN > 8), before proceeding with downstream applications.

### Semi-quantitative and quantitative PCR

RNA was retrotranscribed into cDNA using the iScript™ cDNA synthesis kit (BioRad, 1708891). PCR was performed using Phusion Universal qPCR Kit (Life Tech, F566L). PCR products were loaded in a 2% agarose gels, supplemented with Atlas DNA stain and separated by standard gel electrophoresis. DNA gels images were acquired with an UV scanner (UVITEC). Real-time quantitative PCR was performed with qPCRBIO SyGreen Mix (PCRBiosystems, PB20.14-05), according to the manufacturer instructions, and the CFX96 Real Time PCR thermocycler (Bio-Rad). The data were processed using Bio-Rad CFX Manager software (v.3.1), while gene expression quantification and statistical analyses were performed with GraphPad PRISM (v.6.01). The following primers were used to evaluate the expression of *Foxa1*, Fw: 5’-CATGAGAGCAACGACTGGAA-3’ and Rev: 5’- TTGGCGTAGGACATGTTGAA-3’; *Tbp*, Fw: 5’-CGGTCGCGTCATTTTCTCCGC-3’ and Rev: 5’-GTGGGGAGGCCAAGCCCTGA-3’; *Gapdh*: Fw: 5’-GAGAGTGTTTCCTCGTCCCG-3’ and Rev: 5’-ACTGTGCCGTTGAATTTGCC-3’.

### ChIP sequencing data analysis

Published ChIP-seq fastq files were retrieved from GEO (GSE128867). Reads were aligned to the mm10 reference genome with bowtie (v1.3) and peak calling was performed with MACS2 v2.2.7 in narrow mode, with parameters “--keep-dup all -m 3 30 --format BAMPE -- pvalue 0.05”. Due to the extremely low number of sequenced reads, we discarded two samples (R219S-FOXA1-rep1 and R219S-AR-rep1) from the dataset. Next, Irreproducible Discovery Rate (IDR v2.0.3) framework was used to select high reproducible peaks between the two replicate samples per condition (except for R219S). Only peaks with an IDR < 0.05 were kept. These remaining regions have been annotated with the R package ChIPseeker v1.28.3 (*59*), with a range to define a promoter peak of ± 2.5 kb. Distal elements were defined as all the non-promoter regions, including Distal Intergenic, UTR, Intronic and Exonic. Finally, the peaks overlap between different condition was performed with the R package ChIPpeakAnno v3.32 (*60*). bamCoverage from deepTools v3.5.1(*61*) was used to create BigWig files, with parameters “--binSize 50--extendReads”. BigWigs from the two replicates were merged with wiggletools mean (*62*). deepTools functions computeMatrix and plotHeatmap were used to visualize the ChIP-seq signals over the called peaks.

### ChIP-seq and Rna-seq Data Integration

To allow the comparison between gene expression and transcription factor binding data, we selected differentially expressed genes in ENRADA vs ENRAD- from the bulk RNA-seq analysis (|log2FC| ≥ 1 and adj. P-value < 0.05). Both up and down regulated genes were intersected with the target genes from the ChIP-seq. Specifically, each distal or promoter peak was assigned to its nearest gene based on the distance from the promoter region, ensuring a single gene (target) association for each peak. Venn diagrams and heatmaps were created to visualize the overlap. Heatmaps were plotted with the R package ComplexHeatmap (*63*). Moreover, the hyper-geometric test was used to calculate the significance of the intersection between up regulated genes and transcription factor targets from the ChIP-seq. As background for the test, it was set the total number of expressed genes from the RNA-seq (normalized read count across samples ≥ 10).

### RNA sequencing data analysis

cDNA libraries were prepared with TruSeq stranded mRNA library prep Kit (Illumina, RS-122-2101) using 1 μg of total RNA. RNA sequencing was performed on an Illumina HiSeq 2500 Sequencer using standard Rapid Run conditions at the Next-Generation Sequence Facility of University of Trento (Italy). The reads obtained from each sample were on average 25 million, 100 base pairs long, and single-ended. Adapter trimming and quality-base trimming were performed on the FASTQ file generated by the Illumina HiSeq2500 sequencing machine using Trimmomatic-v0.35 (*64*). The reads were aligned to the Mus Musculus genome (mm10) using STAR-v2.6.0 (*65*) with a maximum mismatch of two and default settings for all other parameters. Then, uniquely mapped reads were selected, and individual sample reads were quantified using HTSeq-count v0.5.4 (*66*) tool to obtain gene-level raw counts based on GRCm38.92 Ensembl (www.ensembl.org) annotation. Individual sample counts were normalized via Relative Log Expression (RLE) using DEseq2 (*67*), which was also used to perform differential expression analyses. P-values were adjusted for multiple hypothesis testing using the method of Benjamini and Hochberg. Differentially expressed genes between each comparison were those genes with absolute fold-change > 1 and adjusted p-value < 0.05. Heatmaps were created with the R package ComplexHeatmap (*63*). Specifically, the genes were annotated as indicated in Crowley et. al 2020 and hierarchical clustering was performed on the rows with the complete linkage method. The normalized counts obtained from DESeq2 are shown for each gene and condition. Functional enrichment analysis of rescued genes was performed using Metascape (*68*), using GO biological processes gene sets. All genes in the genome were used as enrichment background. Terms with p-value < 0.01, minimum count of 3, and an enrichment factor > 1.5 were collected and grouped into clusters based on membership similarities. The most statistically significant term within each cluster was chosen to represent the cluster. BigWig files have been generated with deepTools v3.5.1 (*61*) with parameters “--binSize 50 --normalizeUsing RPKM”. The three replicates per condition were merged with wiggletools mean (*62*).

### Single Cell Expression Analysis

Log-2 transformed gene counts were downloaded from the Single-cell atlas of the mouse and human prostate (*8*). These data are available on the Broad Institute Single Cell Portal (https://singlecell.broadinstitute.org/single_cell/study/SCP1080/,SCP1081, SCP1082, SCP1083, SCP1084). The clusters’ annotations for each cell type were retrieved as assigned by Crowley et al. 2020 and used for the bar plot and violin plots generation. Wilcoxon rank test was used for the box plot comparisons.

### Subcellular Fractionation and Western blotting

Organoid cell pellets were lysed in RIPA buffer (50 mM Tris-HCl, pH 7.5, 150 mM NaCl, 1% Triton X-100, 1% sodium deoxycholate, 1% NP-40) supplemented with protease (Halt^TM^ protease inhibitor cocktail, Life Tech, 87786) and phosphatase inhibitors (Phosphatase-Inhibitor Mix II solution, Serva, 3905501). NE-PER Nuclear and Cytoplasmic Extraction Kit (Life Tech, 78833) was used for nuclear/cytoplasmic fractionation according to the manufacturer instructions. Protein concentrations were measured using the BCA Protein Assay Kit (Pierce™ BCA Protein Assay kit, Thermo Fisher Scientific, 23225) and a Tecan Infinite M200 Plate Reader. Proteins were resolved via SDS-PAGE and transferred to polyvinylidene difluoride (PVDF) membrane (Merck, GE10600023) with a wet electroblotting system (Bio-Rad). The membranes were blocked with 5% non-fat dry milk or 5% BSA in TBS-T (50 mM Tris-HCl, pH 7.5, 150 mM NaCl, 0.1% Tween20) for 1 hour at room temperature, then incubated with designated primary antibodies overnight at 4°C. After washing, membranes were incubated with HRP-conjugated secondary antibody for 1 hour at room temperature. ECL LiteAblot plus kit A+B (Euroclone, GEHRPN2235) was used to detect immunoreactive bands with an Alliance LD2 device and software (UVITEC). Primary and secondary antibodies used in this study are provided in the Appendix Table 2.

### Molecular dynamics (MD) simulations

A 3D structural model of mutant FOXA1 (deletion of Phe254 and Glu255 henceforth referred to as FOXA1^F254E255^) complexed to DNA was generated using the model of the wild type FOXA1^WT^ – DNA complex that was published earlier (*38*). This complex was subject to atomistic molecular dynamics (MD) simulations using the same protocols that we had adopted to simulate the complex of FOXA1^WT^ with DNA in a previous study (*38*). The MD simulations were carried out with the pmemd.cuda module of the program Amber18 using the Amber ff14SB force field (*69*) for proteins and the amber force field FF99BSC0 (*70*) for DNA. Three independent MD simulations (assigning different initial velocities) were carried out on the FOXA1^F254E255^ -DNA complex for 100 ns each. To enhance the conformational sampling, the conformations of the FOXA1^WT^ – DNA (taken from the previous study (*38*)) and FOXA1^F254E255^ - DNA complexes at the end of the MD simulations were subjected to accelerated MD (aMD) (*71*) simulations as implemented in Amber18. aMD simulations were performed on both systems using the “dual-boost” version (*72*). For the aMD simulations, the conventional MD simulations mentioned earlier were used to derive the required parameters (EthreshP, alphaP, EthreshD, alphaD). aMD simulations were carried out for 500 ns each. Simulation trajectories were visualized using VMD (*73*) and figures were generated using Pymol (De Lano, W., The PyMOL molecular graphics system. De Lano Scientific: San Carlos CA, USA, 2002).

### Statistical Analysis

Data are represented as mean ± standard deviation (s.d.) of at least three independent biological replicates except when otherwise indicated. Differences were analyzed by Student’s t test or one-way ANOVA with respectively Bonferroni’s and Duncan’s post-hoc corrections, using PRISM 6 (GraphPad V. 6.01). P-values < 0.05 were considered significant.

## Acknowledgments

We thank current and former members of the Lunardi laboratory for experimental support and advice. We are grateful to all the staff at the CIBIO core facilities for technical assistance and support in data acquisition and analysis. Illustrations were created with Inkscape and BioRender.com.

## Funding

This work was supported by The Giovanni Armenise-Harvard Foundation (Career Development Award), Italian Ministry of University and Research (PRIN 20174PLLYN), Associazione Italiana per la Ricerca sul Cancro (AIRC-IG 27893), Lega Italiana Lotta ai Tumori (LILT-Bolzano), Fondazione Trentina per la Ricerca sui Tumori (FTRT), and core funding from the Department CIBIO to A.L.; by the Associazione Italiana per la Ricerca sul Cancro (AIRC MFAG 2017-ID 20621) to A.R.; by Italian Association for Cancer Research (AIRC, MFAG-20344) and Worldwide Cancer Research (23–0321) to F.C, by University of Trento (Starting Grants Young Researchers 2019) to A.A, and by grants from the National Institutes of Health (R01 CA238005 and U01 CA261822) to M.M.S. Individual fellowships were awarded from the Fondazione Umberto Veronesi (FUV 2016) to A.A., (FUV 2016-2017) to F.Ca., from the United States Department of Defence (W81XWH-18-1-0424) to F.Ca. and from the University of Trento (Ph.D. fellowship) to D.D.F., V.F., D.D., and D.B.

## Author contributions

Conceptualization, D.D.F., F.Ca., A.L.; data curation, D.D.F., A.A., A.L.; formal analysis, D.D.F, D.B., F.Ch., F.Ca., A.L.; funding acquisition, A.L.; investigation, D.D.F, A.A., D.B., S.G., E.M., N.A., G.B., D.D., F.Cu., V.F., M.L., F.G., S.K., C.S.V. A.V., F.Ca.; methodology, D.D.F., F.Ca., A.L.; project administration, D.D.F., A.A., S.G., A.L.; resources, M.M.S., A.R., F.Ch., A.L.; software, D.B., D.D., S.K., C.S.V., A.V., A.R., F.Ch.; supervision, F.Ca., A.L.; validation, D.D.F., F.Ca., A.L.; visualization, D.D.F., A.A., D.B., F.Ch., A.L.; writing – original draft, D.D.F., A.A., M.M.S., F.Ch., F.Ca., A.L.

## Competing interests

The authors declare no competing interests.

## Data and materials availability

All unique/stable reagents generated in this study are available from the Lead Contact upon reasonable request or with a completed Material Transfer Agreement. RNA sequencing data have been deposited in the BioProject database under the accession number PRJNA1064118. All other data supporting the findings of this study are available from the corresponding authors upon reasonable request.

---

**Supplementary Figure S1.**
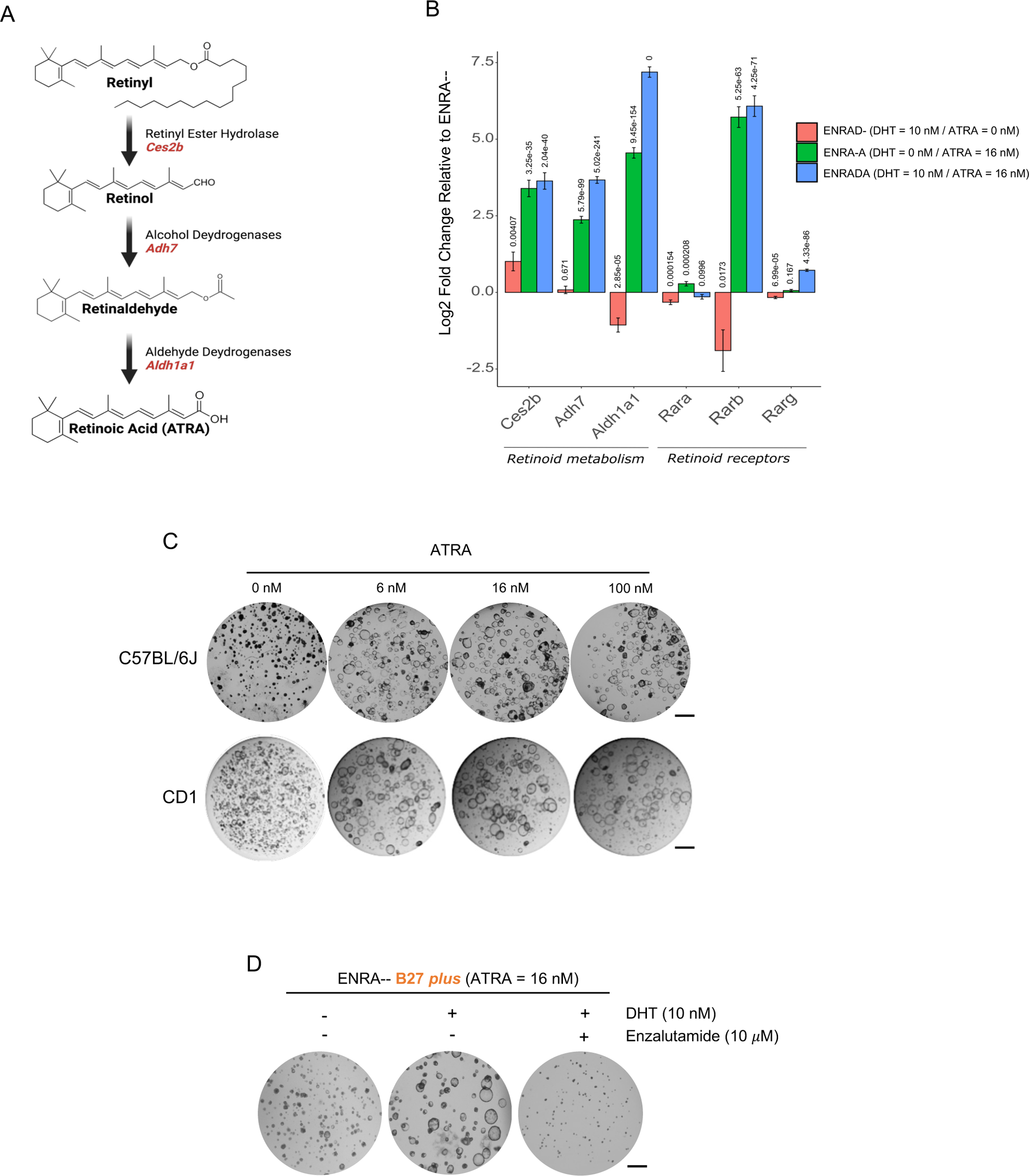
Induction of RA signaling-responsive genes and lumen formation by ATRA and DHT treatment in prostate organoids. (A) Schematic representation of the three main enzymatic steps of retinoid metabolism (made with Biorender). (B) RNA-Seq analysis showing differentially expressed genes involved in the retinoid pathway upon single or combined administration of ATRA and DHT to mPrOs cultured in ENRA-- medium. Data are presented as mean value ± s.d. of n = 3 biological independent replicates. (C) mPrOs (C57BL6/J-upper panel and CD1-lower panel) morphology after 6 days of administration of different concentration of ATRA. Scale bar, 1 mm. N > 3 independent biological replicates. (D) Phenotypic analysis of mPrOs cultured with 16 nM ATRA with or without DHT and Enzalutamide. Scale bar, 1 mm. N = 3 independent biological replicates.

**Supplementary Figure S2.**
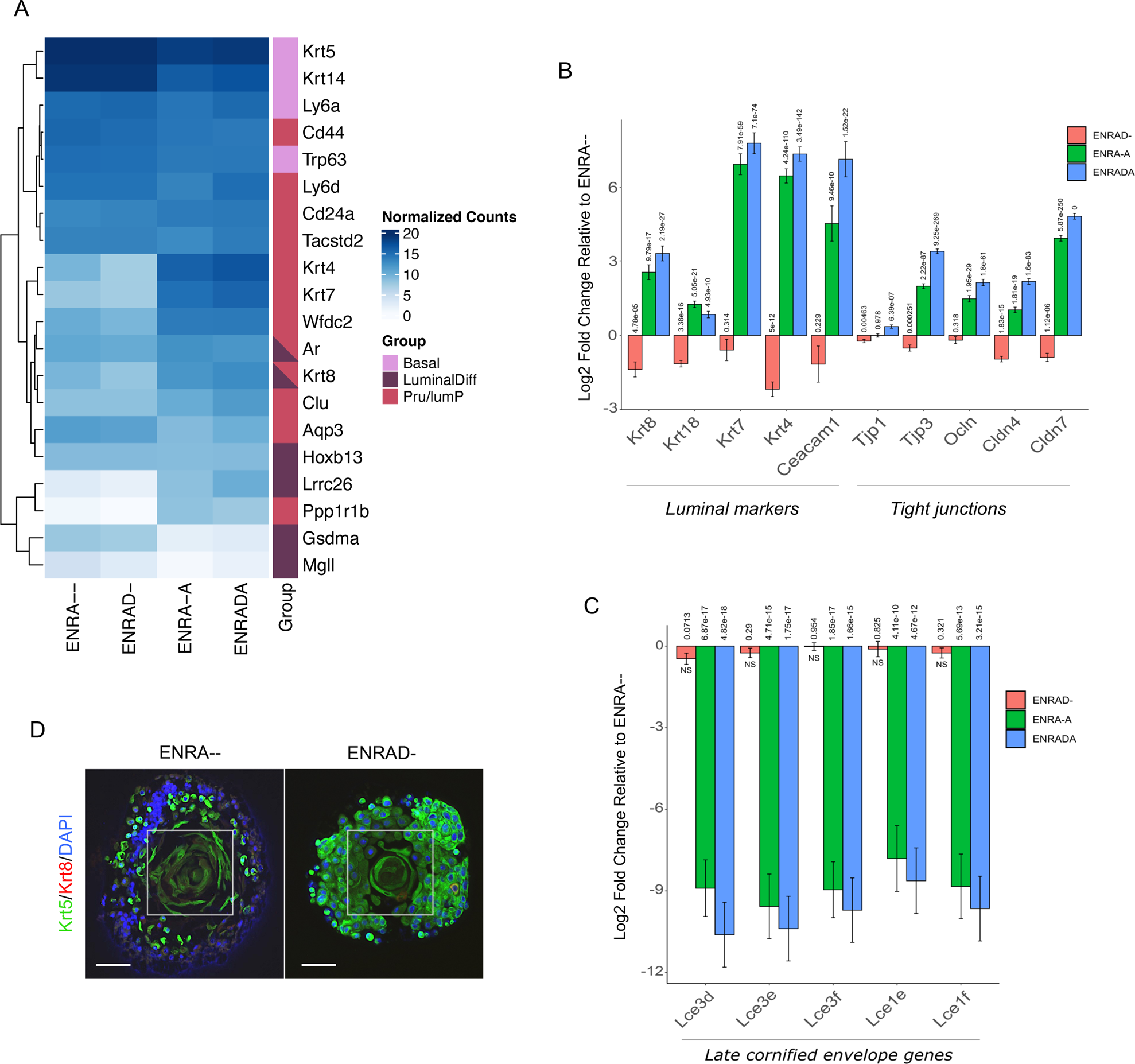
Transcriptional and phenotypic impact of ATRA treatment in prostate organoids. (A) Heatmap showing the expression of a selected panel of genes in mPrOs kept in ENRADA, ENRA-A, ENRAD-, or ENRA-- culture conditions (mean of n = 3 biological independent replicates). Hierarchical clustering with average method has been applied on the heatmap rows. Genes are annotated as basal, luminal differentiated, and periurethral (PrU)/luminal progenitor (LumP) based on Crowley et al. 2020 single cell RNA sequencing analysis. (B) Differential expression of luminal marker and tight-junction genes in mPrOs grown under ENRADA, ENRA-A, ENRAD-, or ENRA-- culture conditions. Data are presented as mean value ± s.d. of n = 3 independent biological replicates. (C) RNA-seq bar plot representation of late cornified envelope genes (*LCE*) the expression of which is robustly repressed by RA signaling. Data are presented as mean value ± s.d. of n = 3 independent biological replicates. (D) Immunofluorescence analysis of Krt5 and Krt8 in mPrOs cultured without DHT and ATRA (ENRA--) or with DHT only (ENRAD). The withe frame marks a peculiar cell morphology noticed only in the absence of ATRA. Scale bars, 100 μm.

**Supplementary Figure S3.**
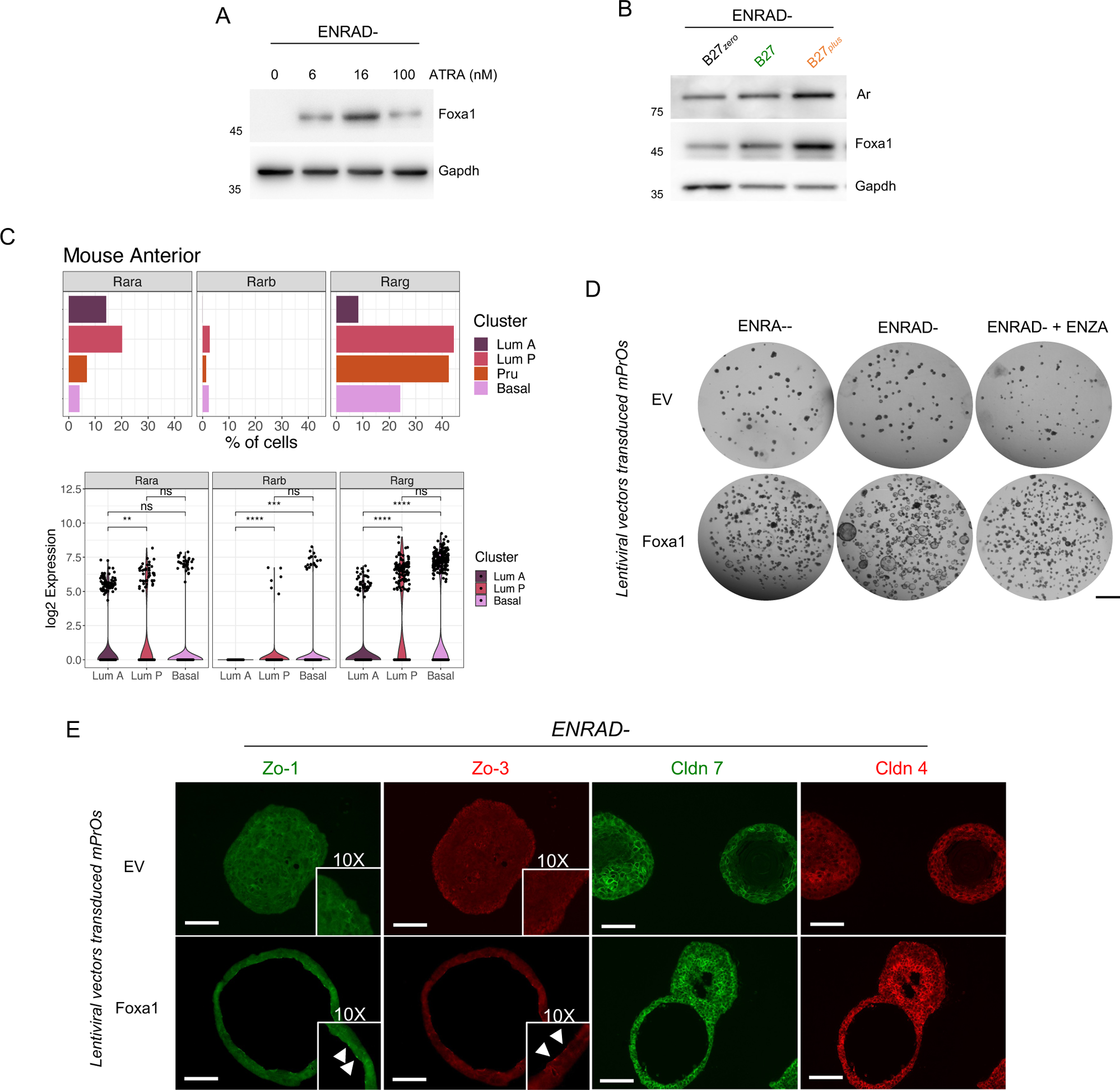
Molecular and phenotypic impact of modulating RA signaling, AR signaling, and Foxa1 expression in prostate organoids. (A) Western blot analysis of endogenous Foxa1 expression in mPrOs treated with different amounts of ATRA. Gapdh is used as loading control. N = 2 independent biological replicates. (B) Western blot analysis of endogenous Ar and Foxa1 expression in mPrOs in presence of different formulations of B27. Gapdh is used as loading control. N = 2 independent biological replicates. (C) Percentage of cells (bar plots) and expression levels (violin plots) of, *Rarα Rarβ*, and *Rarγ* genes in epithelial cell populations of mouse normal prostate (single cell data from Crowley et al., 2020). (D) Morphological analysis of transduced mPrOs ((empty vector (EV) and Foxa1)) cultured without ATRA and DHT (ENRA--), without ATRA with DHT (ENRAD-), and without ATRA with DHT plus Enzalutamide (ENZA 10 μM). (E) Immunofluorescence analysis of Zo-1 (*Tjp1*), Zo-3 (*Tjp3*), Cldn 4, and Cldn 7 expression and localization in transduced mPrOs (EV and Foxa1) cultured without ATRA but with DHT (ENRAD-). Magnification (10x) of Zo-1 and Zo-3 immunostaining are shown to pointing out protein localization. Nuclei are stained with DAPI. Scale bars, 100 μm. N = 2 independent biological replicates.

**Supplementary Figure S4.**
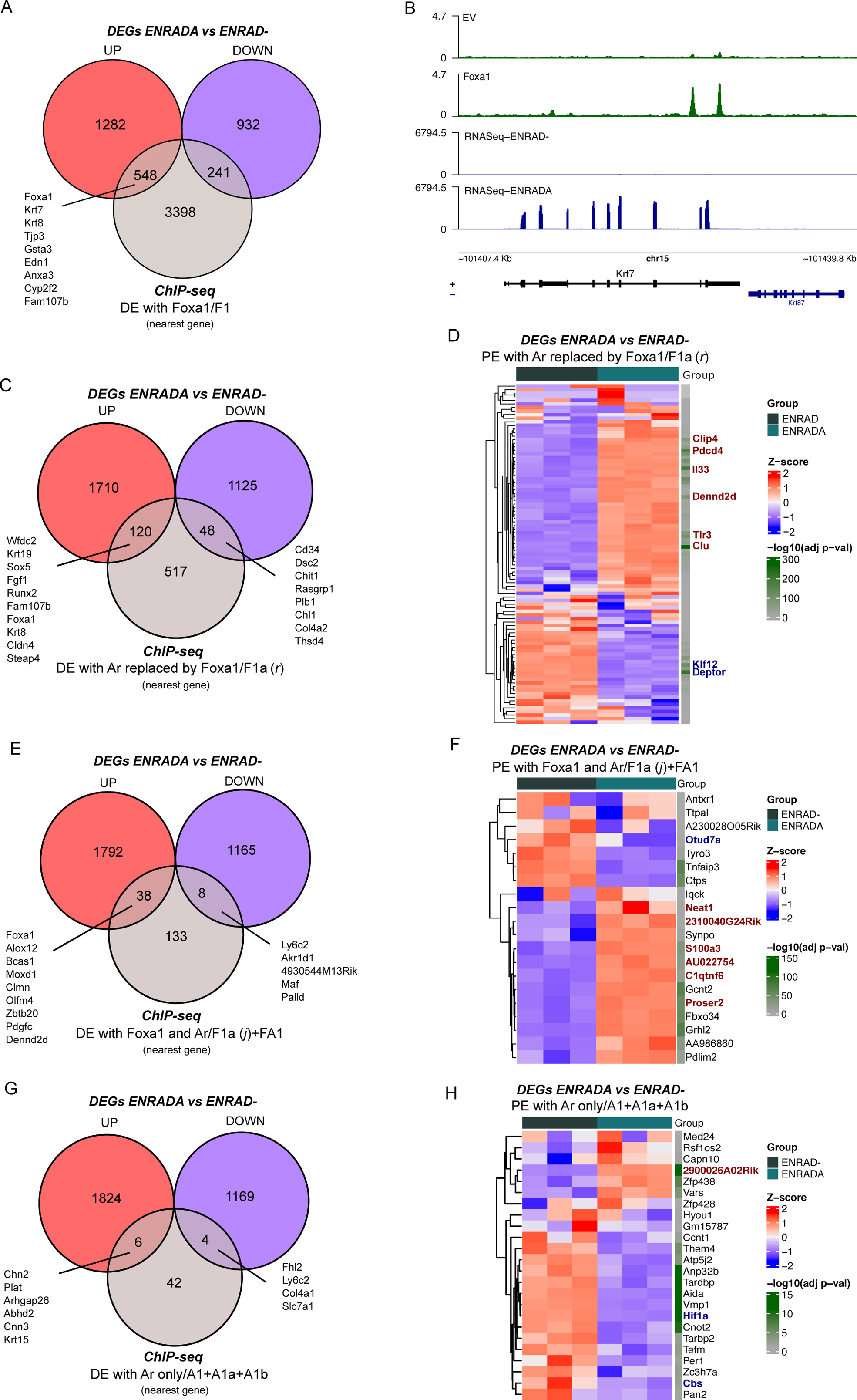
Transcriptional impact of Retinoic acid and Testosterone signaling in prostate organoids. (A) Venn diagrams showing the overlap of differentially expressed genes in mPrOs cultured in ENRADA versus ENRAD, and exogenous Foxa1-bound distal elements (DE (F1)) in the genome. Relevant up-regulated genes in the intersection are highlighted. (B) Genomic snapshot of ChIP-seq (n = 2 pooled replicates) and RNA-seq (n = 3 pooled replicates) signals over the selected gene *Krt7*. (C) Venn diagram showing the overlap between DEGs in mPrOs cultured in ENRADA versus ENRAD and distal elements where exogenous Foxa1 replaces/displaces Ar. Relevant DEGs in the intersections are highlighted. (D) Heatmap showing DEGs in mPrOs cultured in ENRADA versus ENRAD on the promoter of which exogenous Foxa1 displaces/replaces Ar. (E) Venn diagram showing the overlap between differentially expressed genes in mPrOs cultured in ENRADA versus ENRAD and distal elements concomitantly bound by both Ar and exogenous Foxa1. Relevant genes in the intersections are highlighted. (F) Heatmap showing DEGs in mPrOs cultured in ENRADA versus ENRAD whose promoter is concomitantly bound by both Ar and exogenous Foxa1. (G) Venn diagram showing the overlap between DEGs in mPrOs cultured in ENRADA versus ENRAD and distal elements bound by Ar but not by exogenous Foxa1. Relevant genes in the intersections are highlighted. (H) Heatmap showing DEGs in mPrOs cultured in ENRADA versus ENRAD whose promoter is bound by Ar but not by exogenous Foxa1.

**Supplementary Figure S5.**
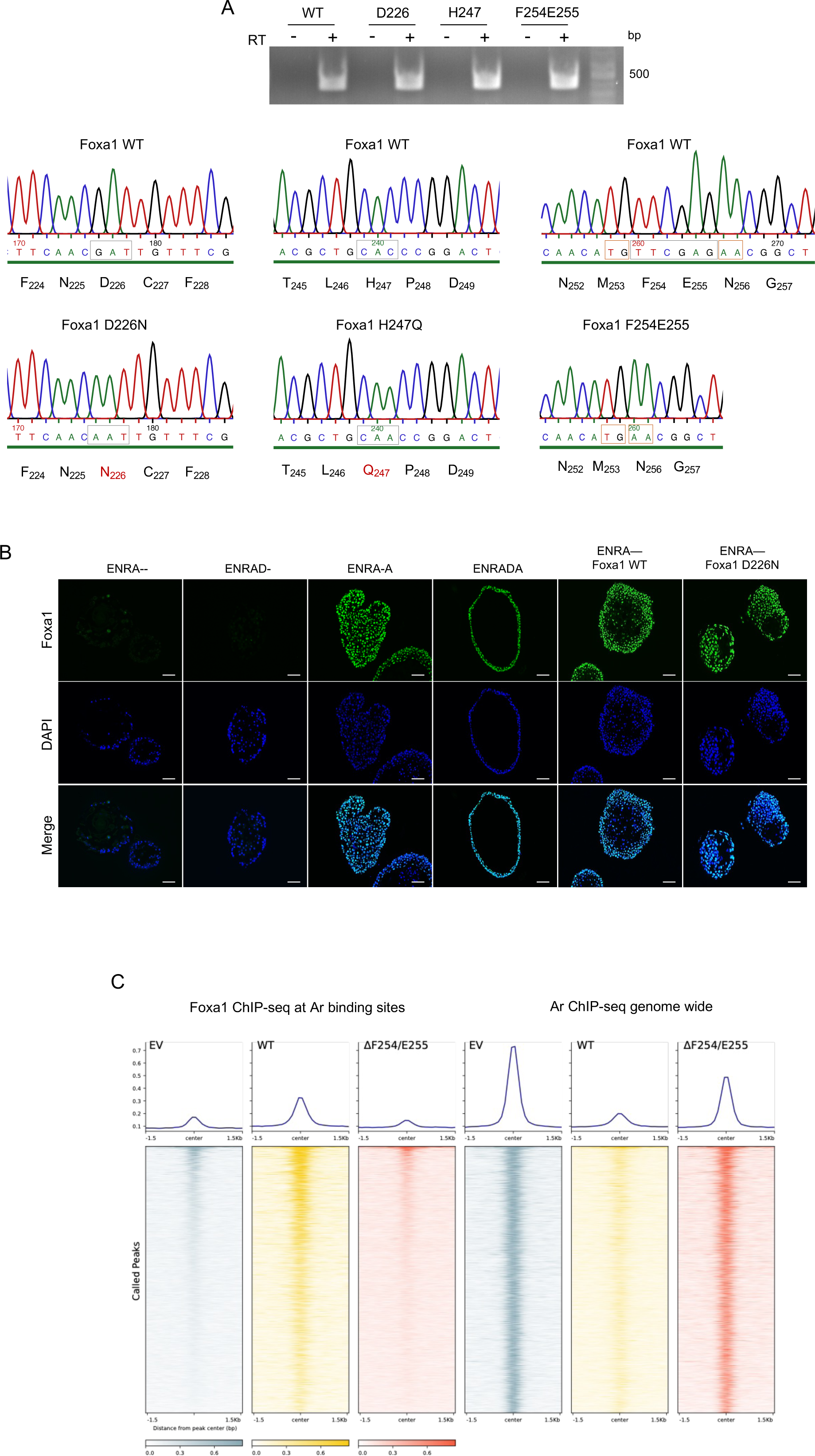
Genetic engineering of prostate organoids with Foxa1 mutant isoforms. (A) RT-PCR and amplicons sequences of wild-type and mutant forms of Foxa1 stably expressed in mPrOs. Spectropherograms highlighting the mutated nucleotides in the different mPrOs lines. (B) Immunofluorescence analysis showing nuclear localization of endogenous and exogenous wild-type and mutant D226N Foxa1 in different growth culture conditions. Scale bar 50 μm. (C) Heatmap showing the signal intensity of Foxa1 and Ar binding over AR genome-wide binding sites (ChIP-seq from Adams et al., 2019). ChIP-seq was performed on mPrOs stably transduced with wild-type Foxa1, Foxa1^F254E255^, or the empty vector (EV) and cultured with DHT but not ATRA.

## Notes

### Competing Interest Statement

The authors have declared no competing interest.

