## Appendix Table 1-2 for "Rarγ -Foxa1 signaling promotes luminal identity in prostate progenitors and is disrupted in prostate cancer"

### Table 1: List of small molecules used in this study

| *Molecule* | *Source* | *Catalog #* |
|---|---|---|
| Egf | PeproTech | 315-09 |
| Noggin | PeproTech | 120-10C |
| A83-01 | Tocris | 2393 |
| DHT | Merck | 10300 |
| ATRA | Merck | R2625 |
| Y-27632 | Merck | Y0503 |
| RARα antagonist (BMS195614) | Cayman | 16029.1 |
| RARβ antagonist (LE135) | Cayman | 14415.1 |
| RARγ antagonist (LY2955303) | Cayman | 25833.1 |
| Pan-RAR antagonist (AGN 193109) | Cayman | 23975.5 |
| Enzalutamide (MDV-3100) | Vinci Biochem | CAY-11596-5 |

### Table 2: List of antibodies used in this study

| *Primary Antibody* | *Source* | *Catalog #* | *Host* | *WB* | *IF* |
|---|---|---|---|---|---|
| Ar | Santa Cruz | sc-816 | Rabbit | 1:1000 | 1:500 |
| Foxa1 | Abcam | ab55178 | Mouse | 1:1000 | |
| Gapdh | ThermoFisher Scientific | MA515738 | Mouse | 1:4000 | |
| β-actin | Merck | A2228 | Mouse | 1:2000 | |
| Cytokeratin 5 | Abcam | ab905901 | Chicken | | 1:500 |
| Cytokeratin 8 | Abcam | ab53280 | Mouse | | 1:500 |
| Zo-1/Tjp1 | Life Tech | 339100 | Mouse | | 1:500 |
| Zo-3/Tjp3 | BioTechne | LS-C313103 | Rabbit | | 1:500 |
| Claudin 4 | Life Tech | 329400 | Mouse | | 1:500 |
| Claudin 7 | Life Tech | 349100 | Rabbit | | 1:500 |
| E-cadherin | Cell Signaling | BK3195T | Rabbit | | 1:500 |
| ***Secondary Antibody*** | ***Source*** | ***Catalog #*** | ***Host*** | ***WB*** | ***IF*** |
| Anti-mouse HRP | Cell Signaling | 7076 | Horse | 1:2000 – 1:8000 | |
| Anti-rabbit HRP | Cell Signaling | 7074 | Goat | 1:2000 – 1:8000 | |
| Mouse Alexa Fluor 488 | LifeTech | A21202 | Donkey | | 1:500 |
| Chicken Alexa Fluor 633 | LifeTech | A21103 | Goat | | 1:500 |
| Rabbit Alexa Fluor 594 | LifeTech | A21207 | Donkey | | 1:500 |
